# High contiguity *de novo* genome sequence assembly of Trifoliate yam (*Dioscorea dumetorum*) using long read sequencing

**DOI:** 10.1101/2020.01.31.928630

**Authors:** Christian Siadjeu, Boas Pucker, Prisca Viehöver, Dirk C. Albach, Bernd Weisshaar

## Abstract

Trifoliate yam (*Dioscorea dumetorum*) is one example of an orphan crop, not traded internationally. Post-harvest hardening of the tubers of this species starts within 24 hours after harvesting and renders the tubers inedible. Genomic resources are required for *D. dumetorum* to improve breeding for non-hardening varieties as well as for other traits. We sequenced the *D. dumetorum* genome and generated the corresponding annotation. The two haplophases of this highly heterozygous genome were separated to a large extent. The assembly represents 485 Mbp of the genome with an N50 of over 3.2 Mbp. A total of 35,269 protein-encoding gene models as well as 9,941 non-coding RNA genes were predicted and functional annotations were assigned.

## 1. Introduction

The yam species *Dioscorea dumetorum* (trifoliate yam) belongs to the genus *Dioscorea* comprising about 600 described species. The genus is widely distributed throughout the tropics [1] and includes important root crops that offer staple food for over 300 million people. Eight *Dioscorea* species are commonly consumed in West and Central Africa, of which *D. dumetorum* has the highest nutrient value [2]. Tubers of *D. dumetorum* are protein-rich (9.6%) with a fairly balanced essential amino acids composition [3]. The provitamin A and carotenoid contents of the tubers of deep yellow genotypes are equivalent to those of yellow corn maize lines selected for increased concentrations of provitamin A [4]. The deep yellow yam tubers are used in antidiabetic treatments in Nigeria [5], probably due to the presence of dioscoretine, which is a bioactive compound with hypoglycaemic properties [6]. Yet, *D. dumetorum* constitutes an underutilized and neglected crop species despite its great potential for nutritional, agricultural, and pharmaceutical purposes.

Unlike other yam species, the agricultural value of *D. dumetorum* is limited by post-harvest hardening, which starts within 24 h after harvest and renders tubers inedible. Previous research showed that among 32 *D. dumetorum* cultivars tested, one cultivar was not affected by the hardening phenomenon [7]. This discovery provides a starting point for a breeding program of *D. dumetorum* against the post-harvest hardening phenomenon. *Dioscorea* cultivars are obligate outcrossing plants that display highly heterozygous genomes. Thus, methods of genetic analysis routinely used in inbreeding species such as linkage analysis using the segregation progeny of an F2 generation and recombinant inbred lines are inapplicable to yam [8]. Furthermore, the development of marker-assisted selection requires the establishment of marker assays and dense genetic linkage maps. Thus, access to a complete and well-annotated genome sequence is one essential step towards the implementation of comprehensive genetic, genomic and population genomics approaches for *D. dumetorum* breeding. So far, a genome sequence assembly for *D. rotundata* (Guinea yam) [8] and a reference genetic map for *D. alata* (Greater yam) [9] have been released. However, these two species belong to the same section of *Dioscorea* (*D*. sect. *Enantiophyllum*) but are distant from *D. dumetorum* (*D*. sect. *Lasiophyton*) in phylogenetic analyses [10,11]. They also differ in chromosome number [8,12,13] making it unlikely that genetic maps can be directly transferred to *D. dumetorum*. Here, we report long read sequencing and *de novo* genome sequence assembly of the *D. dumetorum* Ibo sweet 3 cultivar that does not display post-harvest hardening.

## 2. Materials and Methods

### 2.1. Sampling and Sequencing

The *D. dumetorum* accession Ibo sweet 3 that does not display post-harvest hardening had been collected in the South-West region of Cameroon in 2013 [7]. Tubers of this accession were transferred to Oldenburg (Germany) and the corresponding plants were cultivated in a greenhouse at 25°C. The haploid genome size of the Ibo sweet 3 genotype had been estimated to be 322 Mbp through flow cytometry [14].

High molecular weigth DNA was extracted from 1g of leaf tissue using a CTAB-based method modified from [15]. After grinding the sample in liquid nitrogen, the powder was suspended in 5 mL CTAB1 (100 mM Tris-HCl pH 8.0, 20 mM EDTA, 1.4 M NaCl, 2% CTAB, 0.25% PVP) buffer supplemented with 300 µL ß-mercaptoethanol. The suspension was incubated at 75°C for 30 minutes and inverted every five minutes. Next, 5 mL dichloromethane were added and the solutions were mixed by inverting. The sample was centrifuged at 11,200 g at 20°C for 30 minutes. The clear supernatant was mixed with 10 mL CTAB2 (50 mM Tris-HCl pH 8.0, 10 mM EDTA, 1% CTAB, 0.125% PVP) in a new reaction tube by inverting. Next, a centrifugation was performed at 11,200 g at 20°C for 30 minutes. After discarding the supernatant, 1 mL NaCl (1 M) was added to re-suspend the sediment by gently flicking the tube. By adding an equivalent amount of 1mL isopropanol and careful mixing, the DNA was precipitated again and the sample was centrifuged as described above. After washing the sediment with 1 mL of 70% ethanol, 200 µL TE buffer (10 mM Tris pH 8.0, 0.1 mM EDTA) containing 2 mg DNAse-free RNaseA were added. Re-suspension and RNA degradation were achieved by incubation over night at room temperature. DNA quality and quantity were assessed via NanoDrop2000 measurement, agarose gel electrophoresis, and Qubit measurement. The short read eliminator (SRE) kit (Circulomics) was used to enrich long DNA fragments following the suppliers’ instructions. Results were validated via Qubit measurement.

Library preparation was performed with 1 µg of high molecular weight DNA following the SQK-LSK109 protocol (Oxford Nanopore Technologies, ONT). Sequencing was performed on four R9.4.1 flow cells on a GridION. Flow cells were treated with nuclease flush (20 µL DNaseI (NEB) and 380 µL nuclease flush buffer) once the number of active pores dropped below 200, to allow successive sequencing of multiple libraries on an individual flow cell. Live base calling was performed on the GridION by Guppy v3.0 (ONT).

A total of 200 ng high molecular weight gDNA was fragmented by sonication using a Bioruptor (Diagenode) and subsequently used for Illumina library preparation. End-repaired fragments were size selected by AmpureXp Beads (Beckmann-Coulter) to an average size of 650 bp. After A-tailing and adaptor ligation fragments that carry adaptors on both ends were enriched by 8 cycles of PCR (Illumina TruSeq Nano DNA Sample Kit). The final library was quantified using PicoGreen (Quant-iT) on a FLUOstar plate reader (BMG labtech) and quality checked by HS-Chips on a 2100 Bioanalyzer (Agilent Technologies). The PE library was sequenced in 2 x 250 nt mode on an Illumina HiSeq-1500.

### 2.2. Genome assembly and polishing

Genome size prediction was performed with GenomeScope [16], findGSE [17], and gce [18] based on k-mer histograms generated by JellyFish v2 [19] as previously described [20] for different k-mer size values. In addition, MGSE [20] was run on an Illumina read mapping with single copy BUSCOs as reference regions for the haploid coverage calculation. Smudgeplot [21] was run on the same k-mer histograms (also for different k-mer size values) as the genome size estimations to estimate the ploidy.

Canu v1.8 [22] was deployed for the genome assembly. Raw ONT reads were provided as input to Canu for correction and trimming. Subsequently, Canu assembled the genome sequence from the resulting polished reads. The following optimized parameters were used “‘genomeSize = 350m’, ‘corOutCoverage = 200’ ‘correctedErrorRate = 0.12’ batOptions = -dg 3 -db 3 -dr 1 -ca 500 -cp 50’ ‘minReadLength = 10000’ ‘minOverlapLength = 5000’ ‘corMhapFilterThreshold = 0.0000000002’ ‘ovlMerThreshold = 500’ ‘corMhapOptions = --threshold 0.85 –num-hashes 512 –num-min-matches 3 –ordered-sketch-size 1000 –ordered-kmer-size 14 –min-olap-length 5000 –repeat-idf-scale 50’”. The parameters we selected were optimized for the assembly of a heterozygous genome sequence and our data set. The value for the genome size, estimated to be 322 Mbp, was increased to 350 Mbp to increase the number of reads utilized for the assembly process. A total of 66.7 Gbp of ONT reads with an N50 of 23 kbp was used for assembly, correction and trimming.

ONT reads were mapped back to the assembled sequence with minimap2 v2.17 [23], using the settings recommended for ONT reads. Next, the contigs were polished with racon v.1.4.7 [24] with -m 8 -x -6 -g -8 as recommended prior to the polishing step with medaka. Two runs of medaka v.0.10.0 (https://github.com/nanoporetech/medaka) polishing were performed with default parameters (-m r941_min_high) using ONT reads. Illumina short reads were aligned to the medaka consensus sequence using BWA-MEM v. 0.7.17 [25]. This alignment was subjected to Pilon v1.23 [26] for final polishing in three iterative rounds with default parameters for the correction of all variant types and –mindepth 4.

Downstream processing was based on a previously described workflow [27] and performed by customized Python scripts for purging of contigs shorter than 100 kbp and calculation of assembly statistics (https://github.com/bpucker/yam). In general, sequences were kept if matching a white list (*D. rotundata*) and discarded if matching a black list (bacterial/fungal genome sequences). Sequences with perfect matches against the genome sequences of plants that were sequenced in the lab in parallel (*A. thaliana, Beta vulgaris*, and *Vitis vinifera*) were discarded as well. Contigs with less than 3-fold average coverage in an Illumina short read mapping were compared against nt via BLASTn with an e-value cut-off at 10^−10^ to identify and remove additional bacterial and fungal sequences.

For the ordering (“scaffolding” according to linkage groups) the *D. dumetorum* assembly we employed *D. rotundata* pseudochromosomes. *D. rotundata* pseudochromosome sequences longer than 100 kbp were split into chunks of 1000 bp and subject to a BLASTn search against the *D. dumetorum* assembly with a word size of 12. Hits were considered if the similarity was at least 70% and if at least 70% of the query length were covered by the alignment. To avoid ambiguous hits against close paralogs or between repeat units, BLAST hits were exclude if the second hit exceeds 90% of the score of the top hit. The known order of all chunks on the *D. rotundata* sequence was considered as a “pseudo genetic map” to arrange the *D. dumetorum* contigs via ALLMAPS v0.9.14 [28].

### 2.3. Genome sequence annotation

Hints for gene prediction were generated by aligning *D. rotundata* transcript sequences (TDr96 v1.0) [8] as previously described [29]. BUSCO v3 [30] was applied to generate a species-specific set of AUGUSTUS gene prediction parameter files. For comparison of annotation results, the *D. rotundata* genome assembly GCA_002260605.1 [8] was retrieved from NCBI. Gene prediction hints of *D. dumetorum* and dedicated parameters were subjected to AUGUSTUS v.3.3 [31] for gene prediction with previously described settings [29]. Various approaches involving AUGUSTUS parameter files for rice and maize genome sequences provided by AUGUSTUS as well as running the gene prediction on a sequence with repeats masked by RepeatMasker v4.0.8 [32] with default parameters were evaluated. BUSCO was applied repeatedly to assess the completeness of the gene predictions. The best results for *D. dumetorum* genome sequence annotation were obtained by using an unmasked assembly sequence and by applying yam specific AUGUSTUS gene prediction parameter files generated via BUSCO as previously described [30,33]. Predicted genes were filtered based on sequence similarity to entries in several databases (UniProt/SwissProt, Araport11, *Brachypodium distachyon* v3.0, *Elaeis guineensis* v5.1, GCF_000005425.2, GCF_000413155.1, *Musa acuminata* Pahang v2). Predicted peptide sequences were compared to these databases via BLASTp [34] using an e-value cut-off of 10^−5^. Scores of resulting BLASTp hits were normalized to the score when searched against the set of predicted peptides. Only predicted sequences with at least 0.25 score ratio and 0.25 query length covered by the best alignment were kept. Representative transcript and peptide sequences were identified per gene to encode the longest possible peptide as previously established [29,35]. GO terms were assigned via InterProScan5 [36]. Reciprocal best BLAST hits against Araport11 [35] were identified based on a previously developed script [27]. Remaining sequences were annotated via best BLAST hits against Araport11 with an e-value cut-off at 0.0001. The Araport11 annotation was transferred to predicted sequences.

Prediction of non-protein coding RNA genes like tRNA and rRNA genes was performed based on tRNAscan-SE v2.0.3 [37,38] and INFERNAL (cmscan) v1.1.2 [39] based on Rfam13 [40].

RepeatModeler v2 [41] was deployed with default settings for the identification of repeat family consensus sequences.

### 2.4. Assembly and annotation assessment

The percentage of phased and merged regions in the genome was assessed with the focus on predicted genes. Based on Illumina and ONT read mappings, the average coverage depth per gene was calculated. The distribution of these average values per gene allowed the classification of genes as phased (haploid read depth) or merged (diploid read depth). As previous studies revealed that Illumina short reads have a higher resolution for such coverage analysis [42], we focused on the Illumina read data set for these analyses. Sequence variants were detected based on this read mapping as previously described [43]. The number of heterozygous variants per gene was calculated and compared between the groups of putatively phased and merged genes. Predicted peptide sequences were compared against the annotation of other species including *A. thaliana* and *D. rotundata* via OrthoFinder v2 [44].

Sequence reads and assembled sequences are available at ENA under the project ID ERP118030 (see File S1 for details). The assembly described in this manuscript is available under GCA_902712375. Additional annotation files including the contigs assigned to organelle genomes are available as a data publication from the institutional repository of Bielefeld University at https://doi.org/10.4119/unibi/2941469.

Alleles covered by the fraction of phase-separated gene models were matched based on reciprocal best BLAST hits of the coding sequences (CDSs) following a previously described approach [27]. Alleles were considered a valid pair that represents a single gene if the second best match displayed 99% or less of the score of the best match. A customized Python script for this allele assignment is available on github (https://github.com/bpucker/yam).

## 3. Results

In total, we generated 66.7 Gbp of ONT reads data representing respectively about 218x coverage of the estimated 322 Mbp haploid *D. dumetorum* genome. Read length N50 of the raw ONT data set was 23 kbp and increased to 38 kbp through correction, trimming, and filtering. Additionally, 13 Gbp of Illumina short read data (about 40x coverage) were generated. After all polishing steps, the final assembly represents 485 Mbp of the highly heterozygous *D. dumetorum* genome with an N50 of 3.2 Mbp (Table 1). Substantial improvement of the initial assembly through various polishing steps was indicated by the increasing number of recovered BUSCOs (File S2). The final assembly displayed more BUSCOs (92.30% out of 1440 included in the embryophyta data set, see File S2 for details on the various BUSCO classes) compared to the publicly available genome sequence assembly of *D. rotundata* (v0.1) for that we detected 81.70% BUSCOs with identical parameters. Since there is no genetic map available for *D. dumetorum*, we transferred linkage group assignments from *D. rotundata* to our assembly. In total, 206 contigs comprising 330 Mbp were assigned to a linkage group, while 718 contigs remained unplaced with a total sequence of 155 Mbp (File S3). One plastid and six mitochondrial contigs were identified based on sequence similarity to *D. rotundata* organelle genome sequences (see https://doi.org/10.4119/unibi/2941469); the assignment was confirmed by very high coverage in the read mapping. Our *D. dumetorum* plastid sequence turned out to almost identical to the data recently provided for the *D. dumetorum* plastome [11].

**Table 1.**
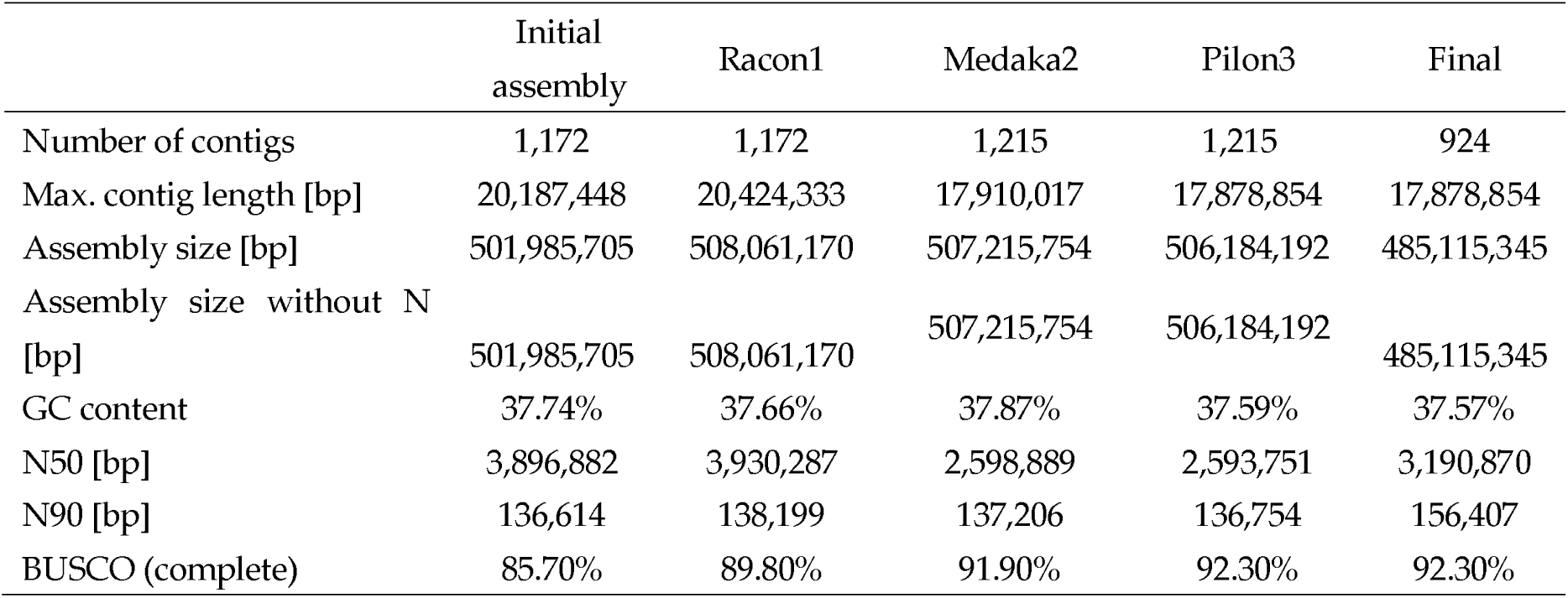
Statistics of selected versions of the *D. dumetorum* genome assembly (see File S5 for a full table).

Haploid genome size estimations based on k-mer distributions of the Illumina sequence reads ranged from 215 Mbp (gce) over 254 Mbp (GenomeScope) to 350 Mbp (findGSE, MGSE) (File S4). The differences between the estimates might be influenced by the repeat content of the *D. dumetorum* genome (see below).

Different gene prediction approaches were evaluated (File S6) leading to a final set of 35,269 protein-encoding gene models. The average gene model spans 4.3 kbp, comprises 6 exons and encodes 455 amino acids (see File S6 for details). The gene prediction dataset for *D. dumetorum* is further supported by the identification of 6,475 single copy orthologs between *D. dumetorum* and *D. rotundata* as well as additional orthogroups (File S7). Based on these single copy orthologs, the similarity of *D. dumetorum* and *D. rotundata* sequences was determined to be mostly above 80% (File S8). If the phase separated allelic gene models were considered (Figure 1), 3,352 additional single copy orthologs were detected. Functional annotation was assigned to 23,835 genes (File S9). Additionally, 9,941 non-coding RNA gene models were predicted including 784 putative tRNA genes (see https://doi.org/10.4119/unibi/2941469). Finally and in addition to gene models encoding proteins and various RNA types, we identified 1,129 repeat consensus sequences with a combined length of 1.3 Mbp (File S10). The maximal repeat consensus length is 17.4 kbp, while the N50 is only 2.5 kbp.

**Figure 1.**
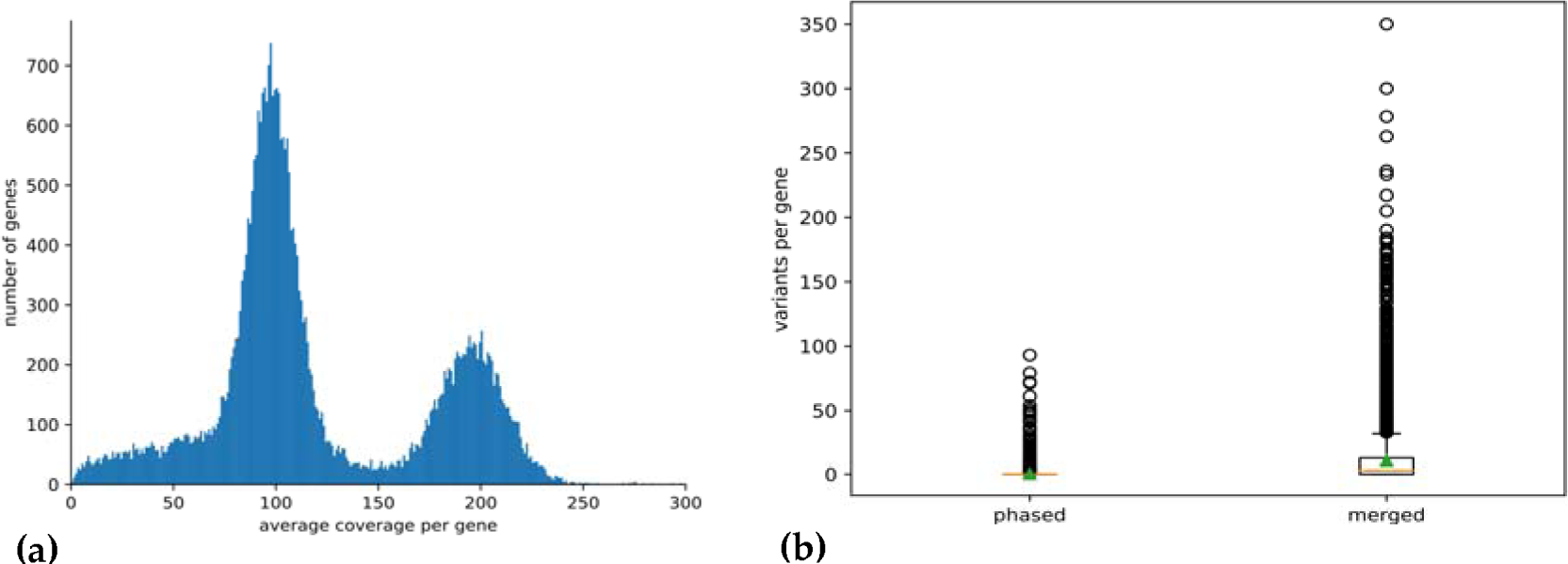
**(a)** Distribution of the average sequencing read depth per gene models. Predicted gene models were classified into phase separated and merged based on the average read depth value deduced from the analysis presented here. The haploid read depth with Illumina short reads ranges from 50-fold to 150-fold. **(b)** Number of heterozygous sequence variants in phase separated and merged genes. The high proportion of heterozygous variants in merged gene models is due to the mapping of reads originating from two different alleles to the same region of the assembly.

Average read mapping depth per gene was analyzed to distinguish genes annotated in separated haplophases as well as merged sequences, respectively (Figure 1, File S11). About 64% of all predicted protein-encoding gene models were in the expected range of the haploid read mapping depth between 50-fold and 150-fold and about 27% are merged with a read depth between 150-fold and 250-fold. Only 6% of all genes show an average read depth below 50-fold and only 1% show an average coverage higher than 250-fold. It should be noted that the gene models annotated in the phase separated part will cover in general two alleles per gene. A total of 22,885 gene models, representing the 64% in the range of the haploid read mapping depth, were sorted into allelic pairs which was successful for 8,492 genes. The findings presented above can be explained by a diploid genome. An analysis with Smudgeplot indicated hints for a tetraploid genome from analysis with a k-mer size of 19, while the other three investigated k-mer sizes supported a diploid genome (File S12).

## 4. Discussion

The release of genome sequences of many model and crop plants has provided new opportunities for gene identification and studies of genome evolution, both ultimately serving the process of plant breeding [45] by allowing discovery of genes responsible for important agronomic traits and the development of molecular markers associated with these traits. Here, we present the first genome sequence for *Dioscorea dumetorum*, an important crop for Central and Western Africa, and the second genome sequence for the genus. Our assembly offers a great opportunity to understand the evolution of yam and to elucidate some biological constraints inherent to yam including a long growth cycle, poor to non-flowering, polyploidy, vegetative propagation, and a heterozygous genetic background [46]. Yam improvement has been challenging due to these factors preventing the genetic study of important traits in yam [47].

Oxford Nanopore sequencing has proven to be a reliable and affordable technology for sequencing genomes thus replacing Illumina technique for *de novo* genome sequencing due to substantially higher assembly continuity [42,48]. Large fractions of the genome sequence were separated into phases, while regions with lower heterozygosity are merged into one representative sequence. Coverage analysis with Illumina read mapping allowed to classify predicted gene models as ‘phased’ or ‘merged’ based on an average coverage around 100 fold or around 200 fold, respectively. While this distinction is possible at the gene model level, whole contigs cannot be classified this way. Several Mbp long contigs comprise alternating phase separated and merged regions. Therefore, it is likely that the contigs represent a mixture of both haplophases with the risk of switching between phases at each merged region. Since the haplophases cannot be resolved continuously through low heterozygosity regions, purging of contigs to reduce the assembly into a representation of the haploid genome might be advantageous for some applications in the future. The bimodal coverage distribution (Figure 1a) supports the assumption that *D. dumetorum* Ibo sweet 3 has a diploid genome. This is supported by Smudgeplot for three out of 4 k-mer sizes tested while the shortest k-mer size used (19) finds indications for tetraploidy. Since a high ploidy would result in more distinct coverage peaks as observed for a genome with up to pentaploid parts [42], we assume that the genome is diploid. The weak hint for tetraploidy might be due to a whole genome duplication event early in the diversification of the genus. The N50 of 3.2 Mbp is in the expected range for a long read assembly of a highly heterozygous plant species which contains quite some repetitive sequences as others reported similar values before [49]. Due to regions of merged haplophases the total assembly size of 485 Mbp is smaller than expected for a fully phase separated “diploid” genome sequence based on the haploid genome size estimation of 322 Mbp.

We noticed an increase of the number of BUSCOs through several polishing rounds. Initial assemblies of long reads can contain numerous short insertions and deletions as these are the major error type in ONT reads [50]. As a result, the identification of CDSs and deduced open reading frames is hindered through apparent disruptions of some CDS. Through the applied polishing steps, the number of such apparent frame shifts is reduced thus leading to an increase of detected BUSCOs.

*D. dumetorum* has 36 chromosomes [12], so with 924 contigs we are far from chromosome-level resolution but considerably better than the other genome assembly published in the genus, that of *D. rotundata* with 40 chromosomes [8]. Knuth [51] circumscribed *D. dumetorum* and *D. rotundata* in two distant sections *D*. sect. *Lasiophyton* and *D*. sect. *Enantiophlyllum*, respectively. Also, phylogenetically the two species are quite distantly related with a last common ancestor about 30 million years ago [11,52]. Comparing our predicted peptides to the *D. rotundata* peptide set [8], we identified about 9,800 single copy orthologs (6,475 in the whole set of 35,269 gene models plus 3,352 with a relation of one gene in *D. rotundata* and two phase-separated alleles in *D. dumetorum*) which could elucidate the evolutionary history of those species. The total number of predicted protein-encoding gene models was determined to be 35,269, but this number includes two copies of about 11,300 gene models (see Figure 1) as these are represented by two alleles each. The CDS-based pairing we performed detected about 8,500 of the theoretical maximum of 11,300 cases which is a good success rate given the fact that close paralogs and also hemizygous genome regions contribute to the detected number of phase-separated gene models. If phase-separated gene models (alleles) are excluded, a number of about 24,000 genes would result for *D. dumetorum*. This fits to the range detected in other higher plant genomes [53,54]. The BUSCO results support this interpretation with about 40% of BUSCOs that occur with exactly two copies. Therefore, the true number of protein-encoding genes of a haploid *D. dumetorum* (trifoliate yam) genome could be around 25,000, also considering that the BUSCO analysis indicated by 5.8% missing BUSCOs that still a small fraction of the genome sequence is missing. This gene number fits well to gene numbers of higher plants based on all available annotations at NCBI/EBI [54]. The average length of genes and the number of encoded amino acids are in the same range as previously observed for other plant species from diverse taxonomic groups [33,55].

It should be noted that the assignment of *D. dumetorum* sequences to the *D. rotundata* pseudochromosomes and indirectly the respective linkage groups contains the risk of incorrect assignments. However, although *D. rotundata* and *D. dumetorum* are evolutionary separated, *D. rotundata* is the most closely related species with genetic and genomic resources.

Our draft genome has the potential to provide a complete new way to breed in *D. dumetorum*, for example avoiding the post-harvest hardening phenomenon, which begins within 24 h after harvest and makes it necessary to process the tubers within this time to allow consumption [2]. The family Dioscoreaceae consists of more than 800 species [56] and the post-harvest hardening phenomenon has only been reported from *D. dumetorum* [57], outlining the singularity of this species among yam species. We predicted a large number of genes, which will include putative genes controlling the post-harvest hardening on *D. dumetorum* and many useful bioactive compounds detected in this yam species, which is considered the most nutritious and valuable from a phytomedical point of view [58]. Ongoing work will try to identify these genes and polymorphisms for making them available for subsequent breeding.

In summary, we present the first *de novo* nuclear genome sequence assembly of *D. dumetorum* with very good contiguity and partially separated phases. Our assembly has no ambiguous bases with a well applicable protein-encoding gene annotation. This assembly unraveled the genomic structure of *D. dumetorum* to a large extent and will serve as a reference genome sequence for yam breeding by helping to identify and develop molecular markers associated with relevant agronomic traits, and to understand the evolutionary history of *D. dumetorum* and yam species in general.

## Supplementary Materials

The following are available online:

File S1: Sequencing overview with ENA identifiers of runs.

File S2: Results of BUSCO analysis of different assembly versions.

File S3: AGP file describing contig assignment to *D. rotundata* pseudochromosomes.

File S4: Genome size estimation overview using four different tools.

File S5: General statistics of different assembly versions.

File S6: Comparison of results from different gene prediction approaches.

File S7: Orthogroups of predicted peptides of *D. rotundata* and *D. dumetorum*.

File S8: Similarity of *D. dumetorum* and *D. rotundata* based on single copy orthologs.

File S9: Functional annotation of predicted genes in the *D. dumetorum* genome sequence.

File S10: Consensus sequences of repeat elements detected in the *D. dumetorum* genome sequence.

File S11: Average short read mapping coverage of predicted genes in the *D. dumetorum* genome sequence.

File S12: Results from Smudgeplot analyses.

## Author Contributions

CS, BP, DCA, and BW designed the study. CS collected the sample. BP performed DNA extraction, ONT sequencing, and genome assembly. PV performed Illumina sequencing. CS and BP processed the assembly. BP performed gene prediction and evaluation. CS and BP wrote the initial draft. BW and DCA revised the manuscript. All authors read and approved the final version of the manuscript.

## Funding

This research was partly funded by the German Academic Exchange Service (DAAD, No. 57299294) with a fellowship to CS.

## Acknowledgments

We thank the Appropriate Development for Africa foundation (ADAF) for the yam collection in Cameroon and for permission to study the plants in Germany in the framework of our mutual protocol agreement. We also thank the German Society of Botany (DBG) for supporting the research stay of CS in Bielefeld. We acknowledge support for the Article Processing Charge by the Deutsche Forschungsgemeinschaft and the Open Access Publication Fund of Bielefeld University.

## Conflicts of Interest

The authors declare no conflict of interest.

## Notes

#### Summary of Updates

Substantial revision

https://github.com/bpucker/yam

