## Supplementary figures and images for "High contiguity *de novo* genome sequence assembly of Trifoliate yam (*Dioscorea dumetorum*) using long read sequencing"

### File S2

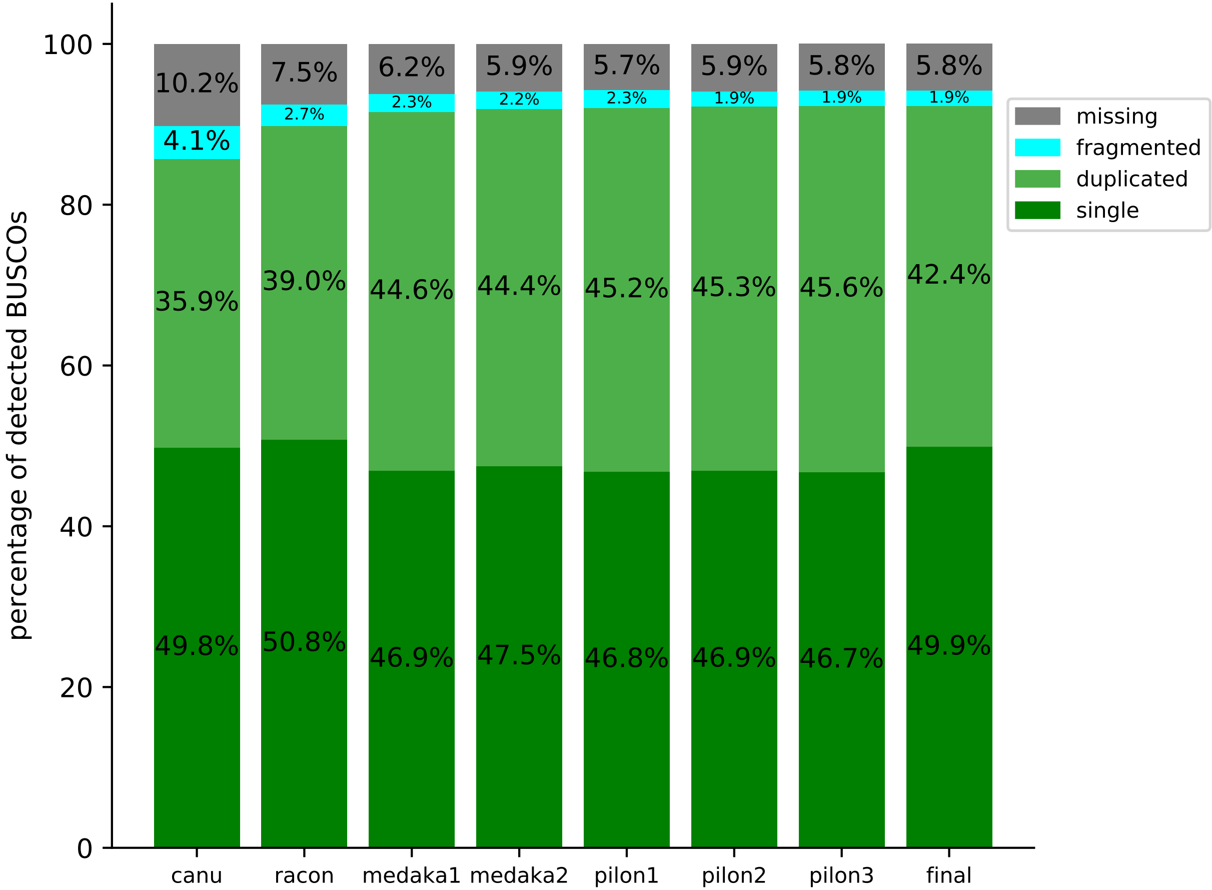

### File S8

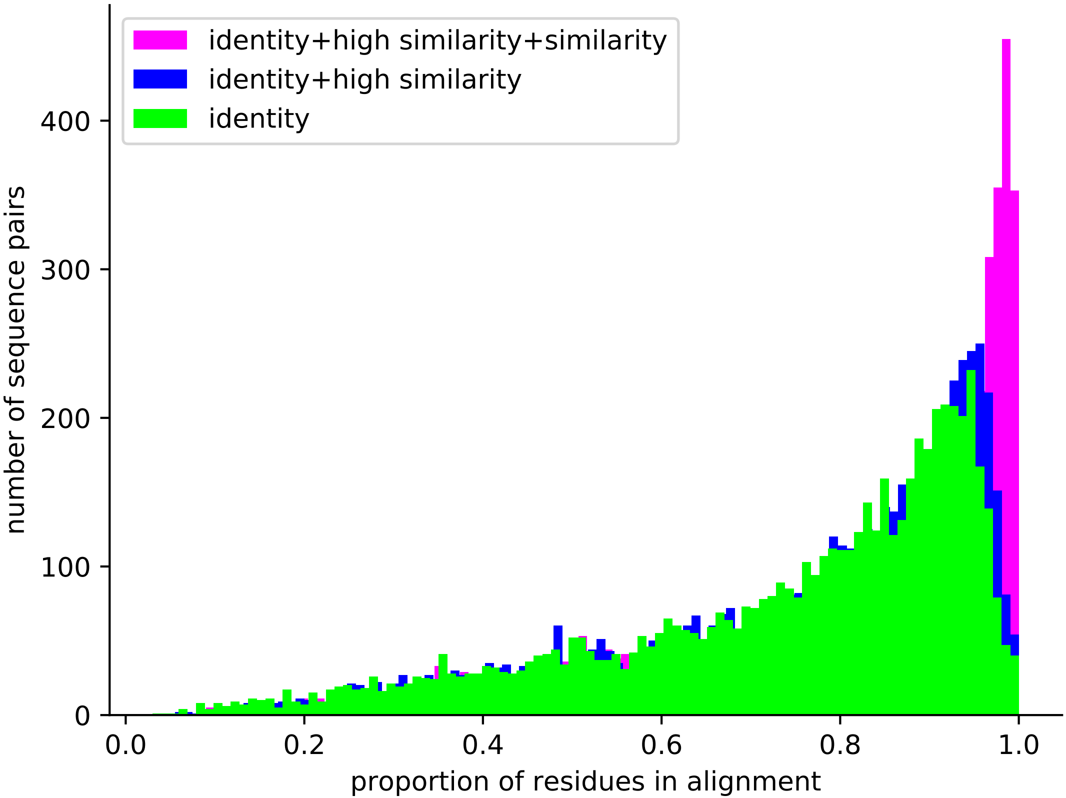

### File S12

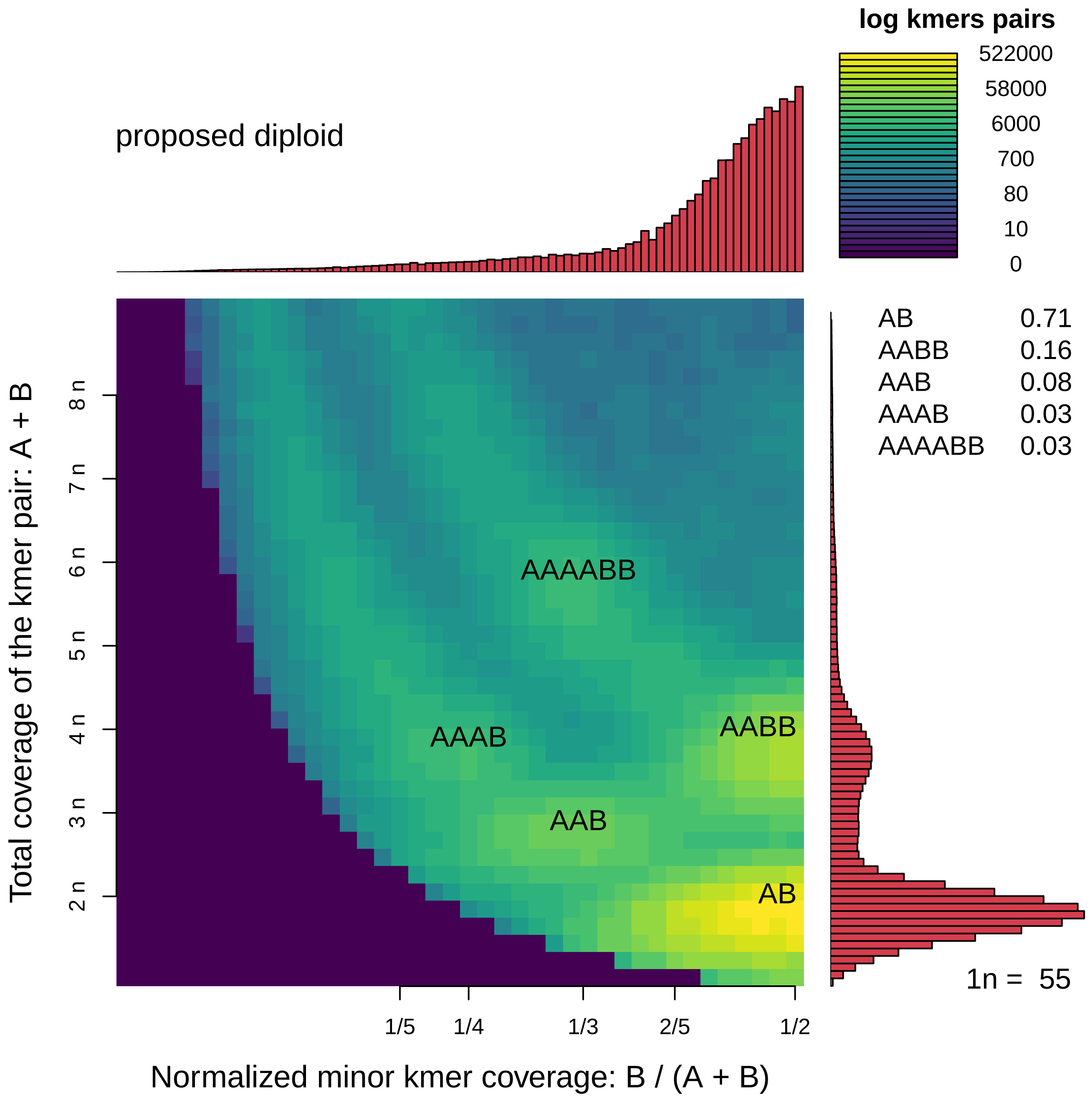
